# Apathy as a Loss of Prior Precision on Action Outcomes

**DOI:** 10.1101/2025.10.06.680685

**Authors:** Rebecca S. Williams, Michelle Naessens, Amirhossein Jafarian, Juliette Lanskey, Frank H Hezemans, Karl Friston, Laura E. Hughes, James B. Rowe

## Abstract

Apathy is common in neurological disease, associated with poor prognosis and limited treatments. Current models posit that goal-directed actions are reduced because costs or effort outweigh the expected reward. We highlight an alternative account of apathy, based on the reduction in precision of prior beliefs about action outcomes. In this preregistered study, we test the hypothesis that precision is encoded in the GABAergic gain of prefrontal superficial pyramidal neurons. Fifty healthy adults undertook a goal-directed task during magnetoencephalography. Estimates of synaptic efficacy or gain were obtained by dynamic causal modelling of induced responses. There was strong evidence of a negative correlation between prior precision and apathy (Bayes Factor=12, p<0.01), and that prior precision was associated with gain in prefrontal and premotor neuronal populations (Posterior probability>0.99). The importance of prior precision and GABAergic gain for goal-directed actions opens new avenues to advance the understanding and treatment of apathy.

## 1. Introduction

Apathy is a common symptom in many neurological and psychiatric conditions, characterised by a reduction of goal-directed actions^1,2^. In lay terms, apathy may be seen as a loss of engagement, reduction in motivation or lack of enthusiasm to engage in everyday activities^3^. Two-thirds of people living with dementia experience at least moderate levels of apathy^4^, with higher prevalence in conditions associated with frontotemporal lobar degeneration^5^. Apathy is associated with higher caregiver burden^6^ and poorer prognosis^7–10^, with limited treatment options^11^. Indeed, apathy may be exacerbated by commonly prescribed medications such as antidepressants^12^. With this in mind, apathy is a priority symptom for treatment^13,14^, but a mechanistic neurocognitive model of apathy is required to support the development of new therapeutics.

Current cognitive perspectives of apathy focus on the cost-benefit analysis of decision-making^15^. Apathetic individuals are proposed to be insensitive to reward^1^, and/or effort avoidant^16^. However, despite successfully modulating reward processing, selective dopaminergic treatments have proven largely ineffective in treating apathy in disease^17,18^. The dopaminergic model of apathy is further challenged by the positive correlation between apathy and impulsivity^19,20^, symptoms traditionally explained by *hypo-* and *hyper*dopaminergic systems respectively. Though such a correlation might be the result of rapidly fluctuating states, this is at odds with clinical observations^20,21^. We instead propose that apathy and impulsivity both represent failures of action-based decision-making, arising from a reduction in the precision of prior beliefs on action outcomes, in the context of the ‘Bayesian brain’^22,23^.

Under the ‘Bayesian brain’ hypothesis, the brain iteratively updates models of the environment (as the cause of sensory inputs) to generate more accurate beliefs about the world, through combining prior knowledge and sensory observations^22,23^. This allows the brain to update expectations in the presence of unexpected changes and/or inform the selection of an appropriate course of action. However, the selection of an appropriate course of action is dependent on predicting the outcome of this action with sufficient precision^24^. For example, imagine sitting at home reading in the evening. The room grows darker as dusk approaches. Most people would turn on the light. For this to happen, they must (A) register the changing light levels, (B) prefer light over dark and (C) believe that the action of switching the light switch will have the desired effect on light levels. A failure in any of these steps would result in an individual sitting longer in the dark room. Failures in step A would result in a world in which one’s prior expectations are never at odds with sensory observations of the environment, meaning there is rarely a sufficient discrepancy between the two to motivate action or passively update a model. It is never “dark enough” to consider turning on the light. Failures in step B can be considered within a reward framework, whereby a lack of action arises due to reward insensitivity. Finally, failures in step C represent a failure of active inference, the focus of this study.

Active inference is the process of undertaking action in order to reduce the discrepancy between prior expectations of the world and sensory outcomes. With insufficient confidence that action can cause the expected outcome (e.g. turning on the light) there is no reason to act at all, even if light is preferred, as the expected consequences of acting and not-acting are too similar: either way the room is expected to remain dark. In short, action initiation and selection is compromised due to uncertainty about the difference in outcome between committing to a course of action or non-action.

We propose that apathy results from such a failure of active inference^25^. Specifically, that apathy is caused by a reduction in prior confidence (cf. precision) on action outcomes, such that there is an insufficient discrepancy between the endpoint of acting and non-acting to motivate engagement. Action therefore becomes *unnecessary* due to a loss of prior precision, rather than being *undesirable* due to a lack of perceived reward or increase in perceived cost. Thereby a loss of prior precision in the prefrontal cortex may lead to apathy, in the same way that weak precision in sensorimotor regions may lead to bradykinesia^26^ and akinesia^27^ (see *figure 1*).

**Figure 1.**
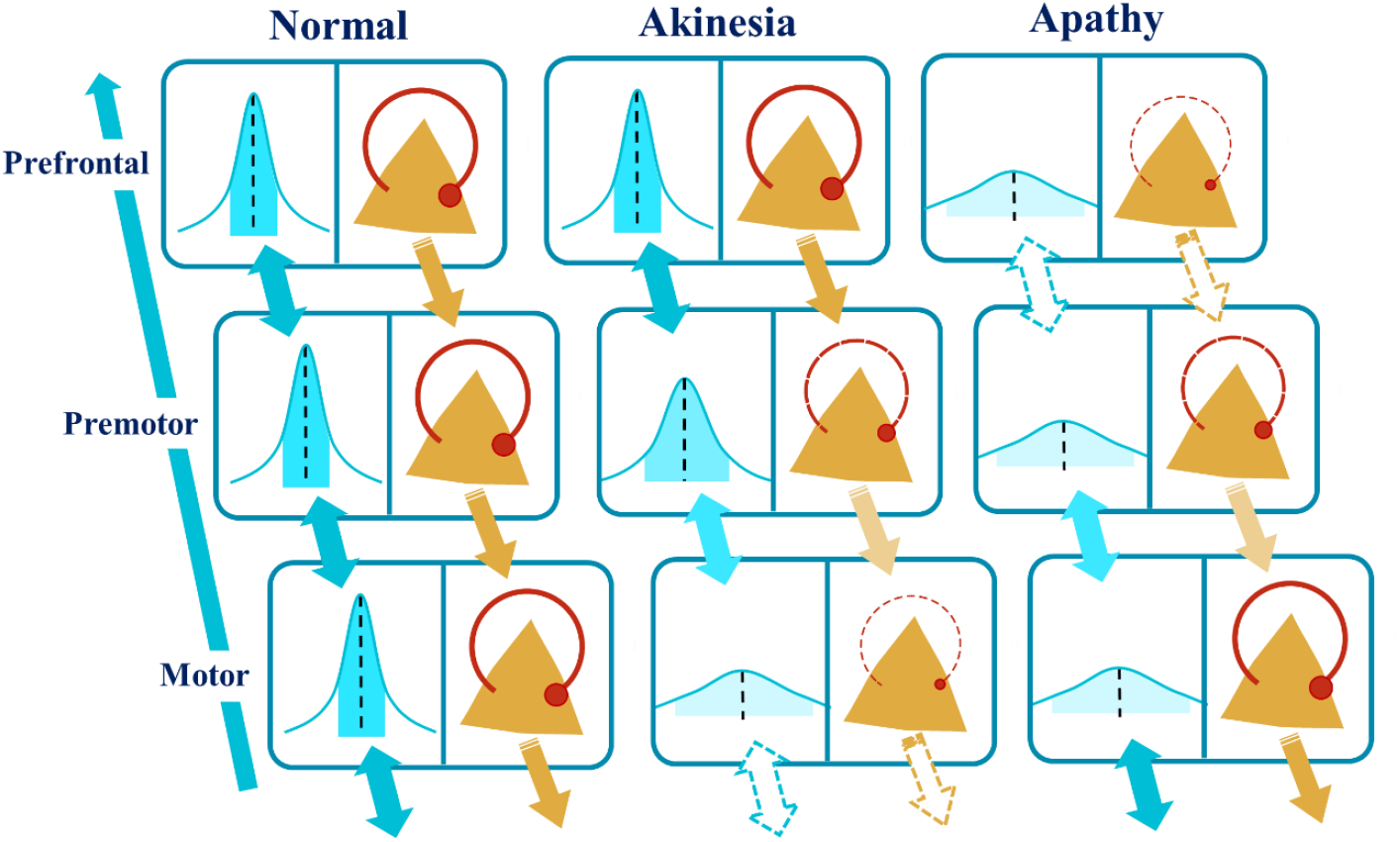
Weaker gain (regulated by GABAergic inhibition) on superficial pyramidal neurons is proposed to lead to apathy or akinesia according to the hierarchical level of the deficit. In the first column, normal priors on action outcomes are communicated effectively across the prefrontal-motor network. However, in the second column a reduction in premotor/motor superficial pyramidal gain results in weak priors on action outcomes at a motoric level, leading to akinesia. In the third column, there is analogous reduction in *prefrontal* superficial pyramidal gain leading to a loss of prior precision at a cognitive level and subsequent apathy.

Theories of the neural determinants of apathy implicate the prefrontal cortex and its subcortical connections, particularly the ventral striatum^28,29^. Both the orbitofrontal and medial prefrontal cortex have been placed high in the decision-making hierarchy in the prefrontal cortex and so may prove key in the communication of action outcomes^24,30^. Both regions have also been associated with apathy and impulsivity^31–33^ with the orbitofrontal cortex playing a central role in updating stimulus value through contextualising sensory input^34^, and the medial prefrontal cortex representing behavioural contexts^35^ crucial to action selection and inhibition^36^. The influence of these regions on action is further mediated through interactions with lateral and medial premotor regions^37^. How then might the reduction of precision in action outcomes be represented and propagated through such a prefrontal-premotor network?

We used dynamic causal models of cortical neurophysiology to determine how prior precision is represented and conveyed across the prefrontal-motor decision-making network. Given the hypothesised role of superficial pyramidal neuron gain in tuning prior precision^38^, we focussed on this neuronal population and their GABAergic regulation. GABA has an established role in tuning oscillatory dynamics^39^ and is depleted in many conditions which feature apathy as a core symptom^40–42^.

Preregistration of the methods, hypotheses, and analyses for this study are available at https://tinyurl.com/3wayrha5. We predicted an association between prior precision of action outcomes and apathy^25^, as assessed using both explicit reporting of beliefs on action outcomes and implicit beliefs extracted from eye gaze. Further, we proposed that intersubject variability in prior precision is associated with variation in pyramidal cell gain in the prefrontal cortex. We tested these hypotheses using behavioural tasks in combination with questionnaires, eye tracking and neuroimaging to confirm and expand on previous apathy studies^25,43^, to open new avenues for research and potential therapeutics.

## 2. Results

We collected data from 50 right-handed, neurologically healthy adult participants who performed a visuomotor ‘goal prior assay’ task during magnetoencephalography (MEG). One additional participant was collected but excluded due to emergence of neurological symptoms during the study. Five datasets were excluded for poor data quality and/or major task instruction deviations with evidence of task misunderstanding^25,43^ (e.g. R^2^ < 0.05).

The remaining 45 participants had a mean age of 64 years (SD=7.26) with mean score of 95.22 on the revised Addenbrookes Cognitive Examination (SD=3.71: ACE-R, max 100), and gender balance of 22:23 (M: F). There was moderate evidence for a correlation between age and prior precision (r=0.37, BF=5.67, p=0.01) such that older individuals had more precise priors. There was no evidence for a correlation between apathy and age (AMI: r=0.06, BF=0.35, p=0.72, CamQUAIT-M: r=-0.05, BF=0.36, p=0.78). There was no evidence for an association between ACE-R score and either prior precision (r=0.29, BF=1.55, p=0.07), or apathy (AMI: r=-0.18, BF=0.62, p=0.25. CamQUAIT-M: r=- 0.08, BF=0.4, p=0.61).

### 2.1 Task Performance

We confirmed the correlation between prior precision and apathy, as indexed by the Apathy Motivation Index total score (AMI; r=-0.42, BF=12.1, p<0.01; *see figure 2A*), and its behavioural subscore (r=- 0.37, BF=5.9, p=0.01; see *figure 2B*). In both cases higher levels of precision were associated with lower apathy scores. These correlations remained significant when factoring in the effect of age on prior precision using partial correlations. There was moderate evidence for a correlation with the AMI social apathy subscore (r =-0.35, BF=3.97, p=0.02) but no evidence for or against a correlation with the AMI emotional apathy subscore (r =-0.16, BF=0.55, p=0.29).

**Figure 2.**
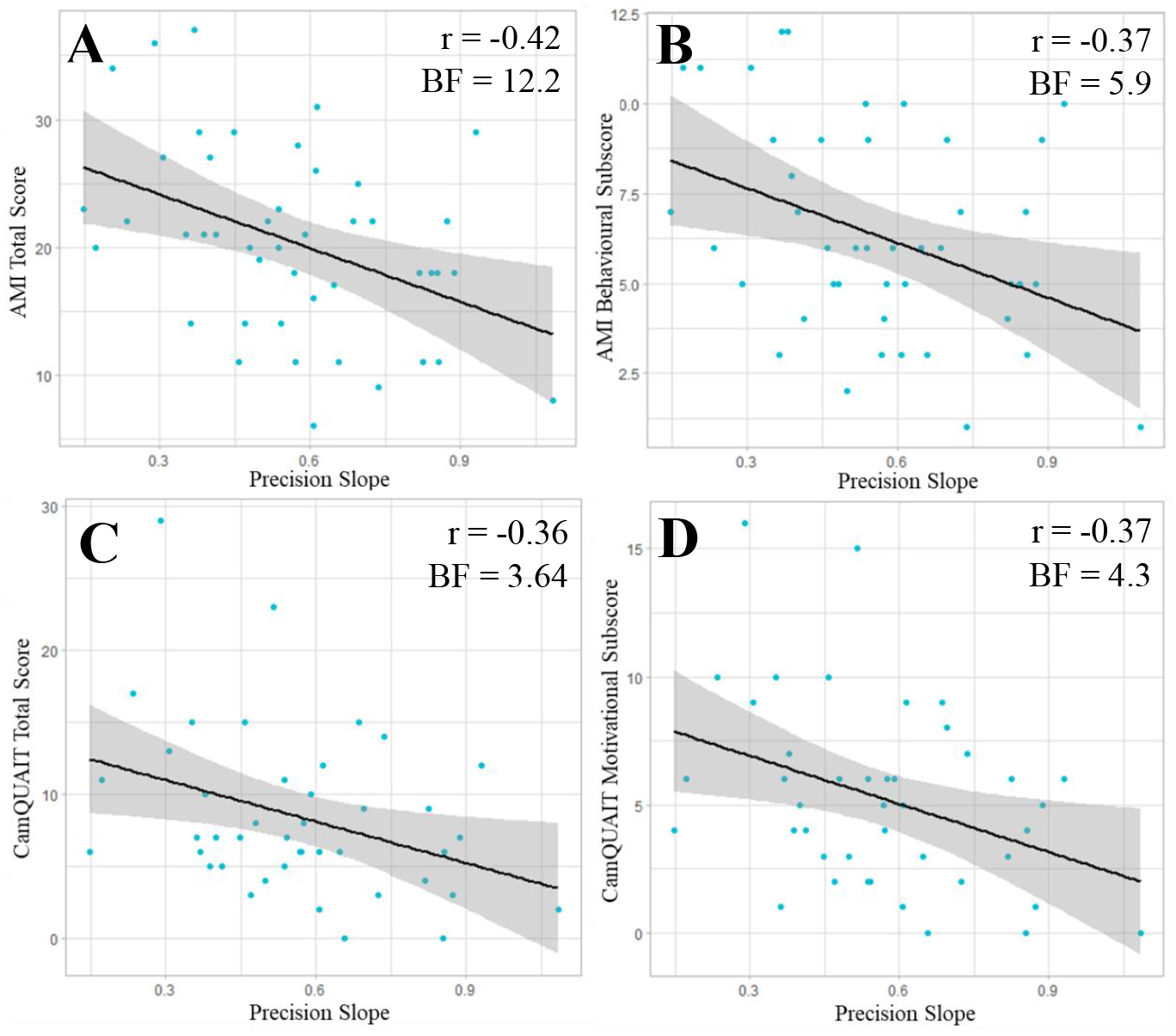
Correlations between prior precision and A) total score on the Apathy Motivation Index (AMI), B) behavioural subscale of the AMI as used in Hezemans et al (2020), C) total score on the Cambridge Questionnaire for Apathy and Impulsivity Traits (CamQUAIT), and D) motivational subscore of the CamQUAIT.

There was evidence for a correlation between prior precision and apathy as indexed by the motivation subscale of the Cambridge Questionnaire for Apathy and Impulsivity Traits (CamQUAIT; r=-0.37, BF=4.34, p=0.02, see *figure 2D*); and the CamQUAIT total score (r=-0.36, BF=3.64, p=0.02; *see figure 2C*). There was no evidence for a correlation with the impulsivity subscale (r=-0.27, BF=1.33, p = 0.08). Moreover, there was no evidence for a correlation between prior precision and A) score on the Barratt Impulsivity Scale (BIS; r=-0.23, BF=0.97, p=0.12) or B) score on the Snaith-Hamilton Pleasure Scale (SHAPS; r=-0.26, BF=1.3, p=0.08). There was moderate evidence for a correlation between total scores on the AMI and CamQUAIT (r=0.39, BF=6.1, p=0.01), primarily driven by the CamQUAIT motivation subscale (r=0.46, BF=21.36, p<0.01) rather than the impulsivity subscale (r=0.22, BF=0.82, p=0.16). There was no evidence for or against a correlation between the total AMI score and BIS (r=0.04, BF=0.34, p=0.82) but there was strong evidence for a correlation between the AMI and SHAPS (r=0.47, BF=40.5, p<0.01).

### 2.2 Eye Tracking

Eye tracking data of adequate quality and calibration were available for 39 participants. As shown in figure 3, participants were able to fixate and anticipate the virtual ball, with gaze along the x axis of the screen, beginning on the ball’s starting position before quickly shifting to fixate around the location of the target after ball release in both non-catch trials (when the ball was visible for the duration of its trajectory) and in catch trials (when the ball disappeared part-way through its trajectory).

**Figure 3.**
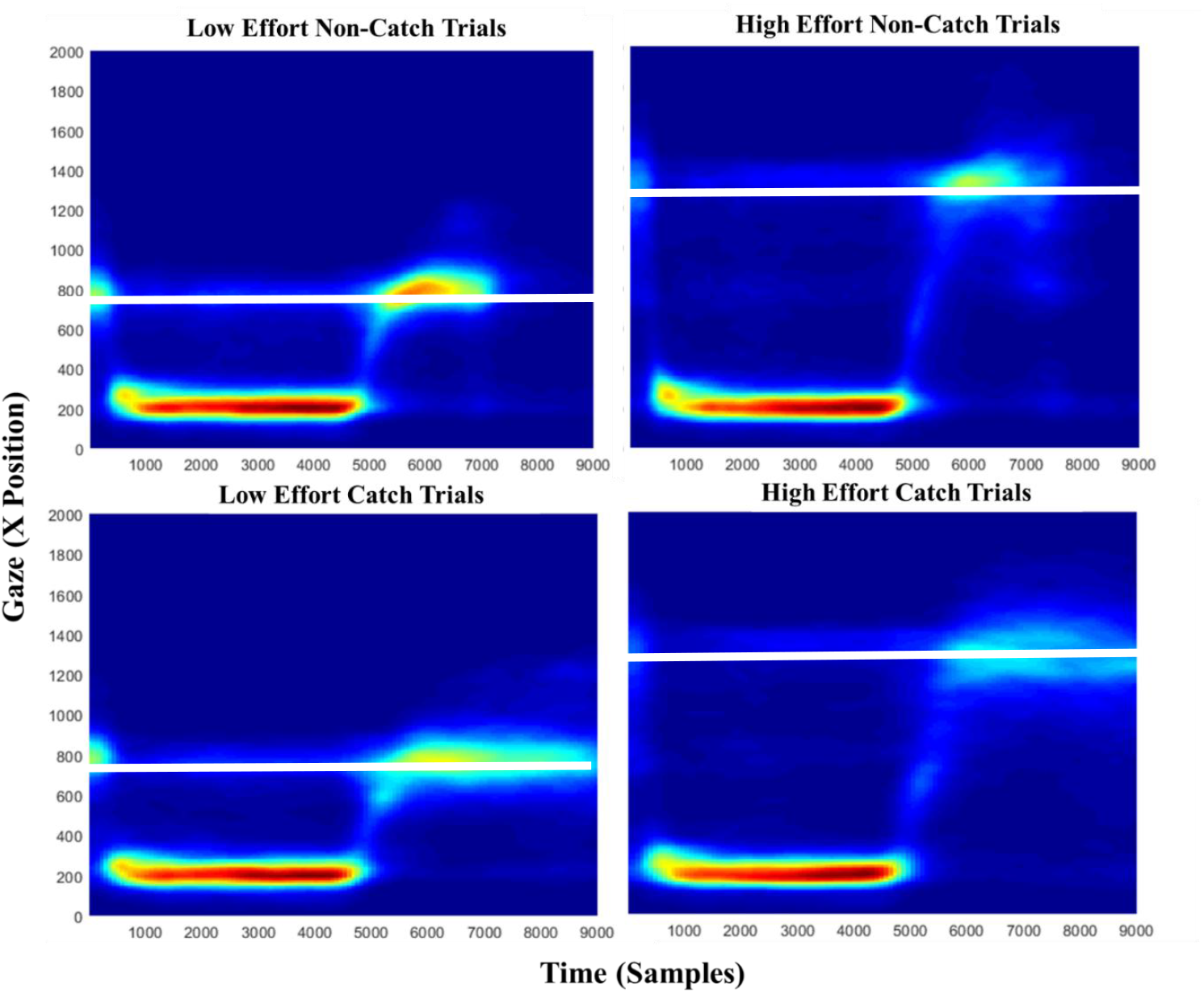
Horizontal gaze position of eye gaze (“X position” on screen) against time (samples, 1 ms) in low and high effort non-catch trials (target visible to participant) and catch trials (target not visible). The horizontal white lines indicate the position of the target at 787 pixels in the low effort trials and 1344 pixels in the high effort trials. Data are averaged epochs across all available trials, and aligned to the start of each trial.

We found very strong evidence for a correlation between true ball location and gaze x position in both low and high effort catch trials (Low effort: r=0.16, BF>100, p<0.01. High effort: r=0.09, BF>100, p<0.01). However, more variance was explained when correlating gaze x position with estimated ball position (Low effort: r=0.37, BF>100, p<0.01. High effort: r=0.3, BF>100, p<0.01).

To explore the potential of replacing explicit estimations of ball position with an implicit eye tracking metric, we generated a new proxy of prior precision calculated as the regression slope of eye-tracking error (i.e. the difference between the final gaze position before the estimation grid appears, and the true location of the ball) on performance error. There was evidence for a positive correlation between regression slopes based on estimation error and eye-tracking error (r=0.42, BF=7.57, p<0.01).

### 2.3 Bayesian Hierarchical Modelling (BHM)

Bayesian Hierarchical Modelling corroborated the regression analyses. We ran the models using both estimation error, based on estimates explicitly given by participants, and eye tracking location. Both analyses favoured a model in which the prior was represented by the location of the target and a small fixed lateral shift compared to a model where the prior was centred on the target voxel (Estimation: ΔWAIC=633; Eye tracking: ΔWAIC=617) or a model in which the prior was centred on true performance (Estimation: ΔWAIC=1331; Eye tracking: ΔWAIC=1026).

Both analyses gave evidence for an effect of effort on prior precision, normalised to subjects actual performance^25^, such that higher effort was associated with lower levels of precision (Estimation: B>100, p<.001; Eye tracking: BF=6.17, p<.001). There was evidence for no effect of reward in the manual task estimation analysis and anecdotal evidence for no effect of reward in the eye-tracking analysis (Estimation: BF=0.21, p=0.11; Eye tracking: BF=0.34, p=0.03; *see figure 4*). Given that “shrinkage” of subject-level estimates skews conclusions at a group-level^44^, the correlations between BHM-estimated prior precision (as extracted from the winning model) and measures of interest are not reported.

**Figure 4.**
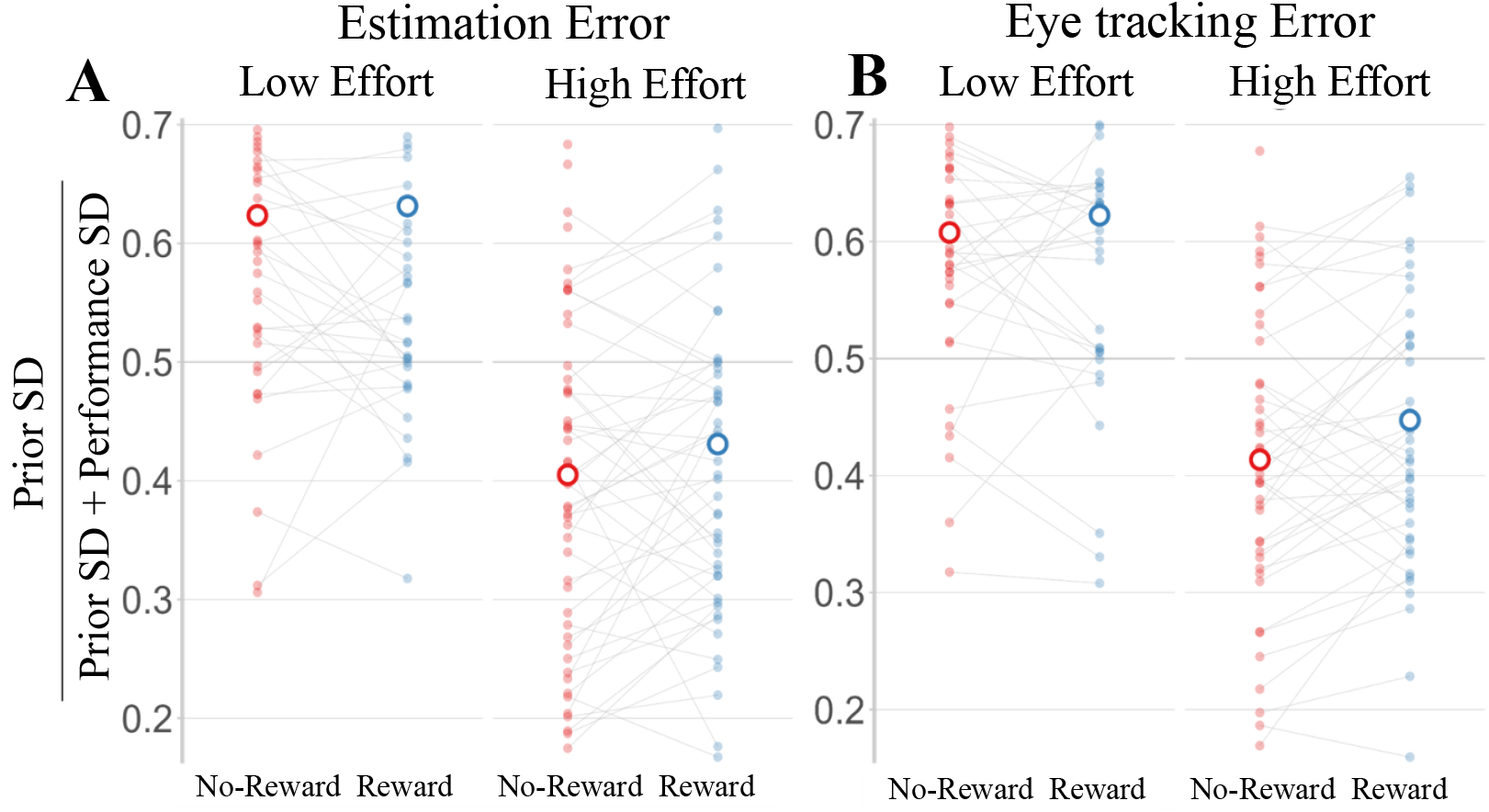
Higher effort increased the prior precision as estimated from Bayesian Hierarchical Modelling of participant actions, whether using (A) explicit estimates given by participants on catch trials or (B) gaze position before the estimation grid appears on screen. Individual estimates of prior SD are illustrated after normalisation to performance SD.

### 2.4 MEG

#### 2.4.1 Spectral Analysis

There was no evidence for or against a correlation between prior precision and beta power in either time window of interest ([-250ms 250ms]: r=-0.26, BF=1.2, p=0.09; [250ms 750ms]: r=-0.02, BF=0.34, p=0.91; see Methods for details). There was anecdotal evidence for a correlation between low beta power and prior precision in the time window around ball stop (r=-0.31, BF=2.27, p=0.04) such that lower levels of prior precision (c.f. higher apathy) were associated with lower beta power. There was no evidence for or against a correlation between high beta power and prior precision ([-250ms 250ms]: r=-0.16, BF=0.54, p=0.3; [250ms 750ms]: r=0.16, BF=0.54, p=0.3). There was no evidence for or against a correlation between measures of apathy and low beta power in the window around ball stop (AMI: r=-0.13, BF=0.47, p=0.39; CamQUAIT-M: r=-0.21, BF=0.72, p=0.2).

#### 2.4.2 Dynamic Causal Modelling

Overall, the dynamic causal model fitted the data well with a median variance explained of 98.3% and median correlation between observed data and the predicted response of our model at 0.996. At a group-level, the inclusion of prior precision as an empirical prior improved model evidence, such that there was strong evidence in favour of the model including prior precision *versus* models excluding prior precision when focussing on all three synaptic parameters of interest: intrinsic connectivity (BF=24.6), extrinsic connectivity (BF=7.8), and neurotransmitter time constants (BF=23.4).

There was very strong evidence that lower prior precision was associated with reduced superficial pyramidal gain in the prefrontal and premotor regions (Pp>0.99), but not in the motor cortex (see *figure 5B*). There was very strong evidence that lower prior precision was also associated with an increased GABA-receptor time constant in the prefrontal and motor cortex (Pp>0.99), but not the premotor region (see *figure 5C*).

**Figure 5.**
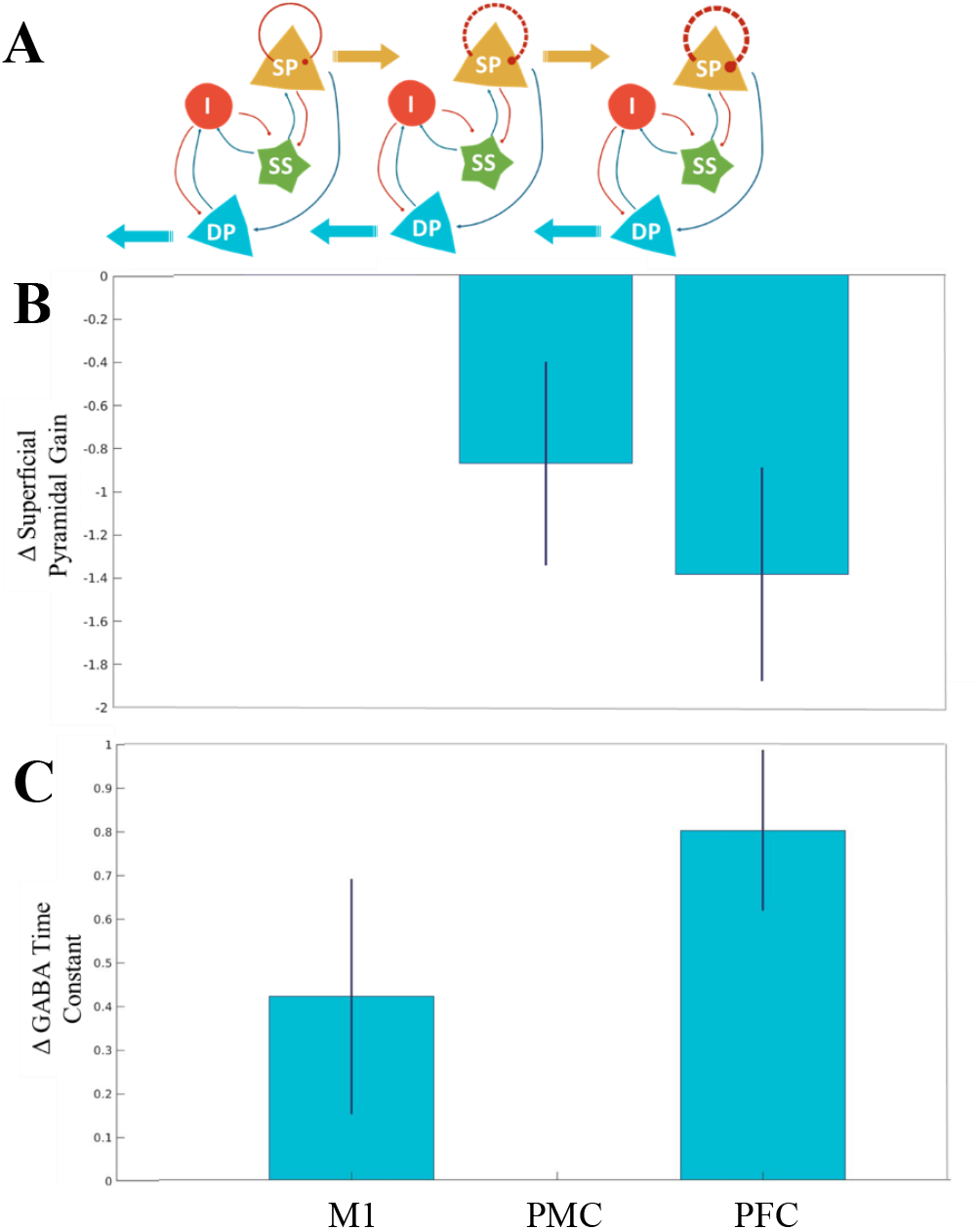
(A) The microcircuit used in each brain source with the superficial pyramidal self-inhibition marked as a red circle. (B) Regions where the superficial pyramidal self-inhibitory connection and (C) regions where the GABAergic rate constant was modulated with a posterior probability >0.99. Neuronal Populations: SP = Superficial Pyramidal I = Inhibitory SS = Spiny Stellates DP = Deep Pyramidal Regions: PFC = Prefrontal Cortex PMC = Premotor Cortex M1 = Primary Motor Cortex

Regarding extrinsic connectivity (between sources), lower prior precision was associated with *reduced* connectivity between the inferior frontal gyrus and premotor area (Pp>0.99). Reduced prior precision was also linked to *higher* levels of connectivity from premotor (Pp>0.95) and motor cortex (Pp>0.99) to the inferior frontal gyrus, and higher connectivity from the premotor to motor cortex (Pp>0.99).

## 3. Discussion

This study provides strong evidence for the association between prior precision and apathy in healthy adults, supporting the hypothesis of apathy as a failure of active inference. Using dynamic causal modelling of cortical physiology, we confirm the association between individuals’ prior precision and GABAergic tuning of superficial pyramidal neurons in prefrontal cortex.

The precision of action-outcomes in the ‘Goal Prior Assay’ task (see *Methods*) correlated with questionnaires’ ratings of apathy, using the Apathy Motivation Index (AMI)^45^, and the motivation subscale of the Cambridge Questionnaire for Apathy and Impulsivity Traits (CamQUAIT-M)^21^. As shown previously^25,43^, this association is primarily driven by the behavioural subscore of the AMI, with weaker correlations to social apathy and no evidence of a correlation with emotional apathy. While active inference related to action-outcomes therefore explains apathy defined as a loss of goal-directed actions, analogous priors of social outcomes or emotional states may explain the components of social and emotional dimensions of apathy^46^.

Dynamic causal modelling of the neurophysiology of the prefrontal-motor hierarchy during the task confirmed that lower levels of prior precision were associated with lower gain on the prefrontal superficial pyramidal neurons and higher GABA time constants. Motor cortical dynamics were less affected. Tuning of the superficial pyramidal neurons in the prefrontal cortex was therefore weaker and slower in individuals with low levels of prior precision. This implies that prior precision reflects efficient communication across the prefrontal-premotor hierarchy. Failures in this system may lead to the propagation of less precise prior expectations from the prefrontal to motor regions, behaviourally resulting in reduced goal-directed action, and therefore increased levels of apathy. This is analogous to the effect of lower-order sensorimotor precision on motoric deficits such as akinesia^27^ and bradykinesia^26^. This hypothesis is in accord with the established role of GABA in tuning oscillatory dynamics^39^ and the direct impact of the superficial pyramidal neurons on the strength of forward and backward connectivity^47^. GABAergic deficits in diseases such as frontotemporal dementia^42^ may therefore explain the high levels of apathy associated with these conditions. Our findings suggest a prefrontal mechanism for apathy, and also the potential to improve apathy by increasing GABA levels with pharmacological treatment

Similar to Hezemans et al (2020)^25^, Bayesian hierarchical modelling demonstrated that prior precision on action outcomes was modulated by effort, but not the low level of reward used in this study (i.e. virtual points). This suggests that low levels of reward are insufficient to overcome deficits in prior tuning, but that reducing perceived effort may be a viable route to behavioural initiation in the presence of low precision. Future work exploring the causal relationship between prior precision and effort avoidance^16^ may offer insight into the potential for behavioural modification as a treatment for apathy given the current lack of consensus regarding the efficacy of non-pharmacological therapies^48^.

This study found strong evidence for a correlation between prior precision when calculated using explicit motor estimations and gaze position. Gaze position was more closely associated with explicit estimates of ball location than with the true location of the ball, suggesting that gaze position may be a suitable proxy for individuals’ estimates of prior precision in future studies. However, if using eye tracking in isolation, it may be necessary to increase the number of trials as we were underpowered to detect a correlation between apathy metrics and prior precision as assessed with eye tracking data alone.

There are several limitations to this study. First, as participants were healthy volunteers in an institutional research panel, apathy levels may be lower than the general population due to volunteer bias. However, there was sufficient variance within our data to power analyses, as evidenced by the Bayes Factors >3. It remains to be seen if these results extrapolate to groups with apathy in the context of clinical disorders, over and above the existing evidence in Parkinson’s disease^25,43^. Second, 10% of datasets were excluded before analysis due to task violations. Eye tracking was a useful tool here to confirm if participants had been responding meaningfully, by comparing their explicit responses with gaze position. However, replacing the estimation grid with an alternative response framework (e.g. moving cursor on screen, touchscreen) may mitigate such problems. Finally, we selected the network for our dynamic causal model based on a combination of prior literature^49^ and source reconstruction of task-based data. We recognise that orbital and medial prefrontal regions are also associated with reward-encoding and apathy in fMRI and morphometric studies^50,51^, but they have very low signal-to-noise in MEG, while the lateral prefrontal region used here is also strongly associated with goal-directed behaviour^52^, reward encoding^53,54^ and volition^55^.

Future studies may include other manipulations of prior precision, either behaviourally or pharmacologically. Given the association with a prefrontal GABA mechanism shown in this study, we predict that GABAergic drugs modulate prior precision and thereby apathy. This is supported by anecdotal reports of the effect of GABA-A agonists on severe amotivational states in small-scale studies^56,57^. Given the impact of noradrenergic drugs on apathy and prior precision^43,58^, generative models could also be used to assess the mechanism of action of noradrenergic treatments.

In summary, we confirm the association between apathy and prior precision on action outcomes. This is underpinned by the tuning of prefrontal prior precision through GABAergic gain of the prefrontal superficial pyramidal neurons. Our results do not challenge the importance of dopamine for representation of the magnitude of rewards, or the importance of reward versus effort in behavioural decisions. However, we suggest that dopaminergic-reward accounts alone are insufficient to explain or treat apathy. Rather, the active inference formulation of goals as prior beliefs raises new approaches to clinical translation and treatment, to improve the prospects for those affected by apathy.

## 4. Methods

Preregistration of methods and analyses for this study are available at https://tinyurl.com/3wayrha5.

### 4.1 Participants

Fifty healthy participants were recruited from the Cognition & Brain Sciences Unit, University of Cambridge, volunteer panel. Participants gave written informed consent, and the study was approved by the local Research Ethics Committee. All experiments were performed in accordance with relevant guidelines and regulations of the University of Cambridge.

Participants were aged 50-85 years old at the time of testing. We chose this age range for our healthy adult participants in order to relate discoveries to future work with people affected by age-related neurodegenerative disorders associated with apathy. Participants had no neurological illness, no history of major psychiatric disorder or seizures, and no current indications of neurodegenerative disease. Participants also had normal or corrected-to-normal visual acuity.

To ensure sufficient power, we employed Bayesian sequential hypothesis testing^59^. We analysed the data after an initial data collection period (N=20) and then subsequently after every 5 participants for the purpose of determining sufficient/insufficient precision for inference (i.e. 3<BF<0.33; agnostic to whether the inference was in favour of the null or alternative hypotheses). Data collection was stopped at N=50 as the preregistered limit.

### 4.2 Procedure

Participants completed two testing sessions. The first session took place in the MEG facility at the Cognition & Brain Sciences Unit, University of Cambridge. The second session entailed 7T MRI acquisition at the Wolfson Brain Imaging Centre, Cambridge University Hospital NHS Trust.

### 4.3 Materials

In the first testing session, participants completed a series of self-report questionnaires including the Apathy Motivation Index (AMI)^60^, Barratt Impulsivity Scale (BIS-11)^61^, and Snaith-Hamilton Pleasure Scale (SHAPS)^62^. The Cambridge Questionnaire for Apathy & Impulsivity Traits (CamQUAIT)^21^ was sent home to be completed by an informant.

Participants then completed the ‘Goal Priors Assay’ task^25^ whilst in an MEG scanner. This task evaluates the mechanisms of apathy in the context of the ‘Bayesian brain’. The goal is to land a virtual ball on a target (see *figure 6*). Participants press a force pad for 3 seconds and the force applied determines the trajectory of the ‘ball’. The ball always travels rightwards and decelerates linearly towards the target. On a pseudo-random subset of trials, the ball disappears, and participants are asked to estimate the end position. They indicate the perceived end position of the ball using a numbered grid on screen. Each participant completed 15 practise trials before 4 blocks of 30 trials, including 10 catch trials, which were varied by reward (points vs. no points) and effort (low force for a close target vs. high force for a far target).

**Figure 6.**
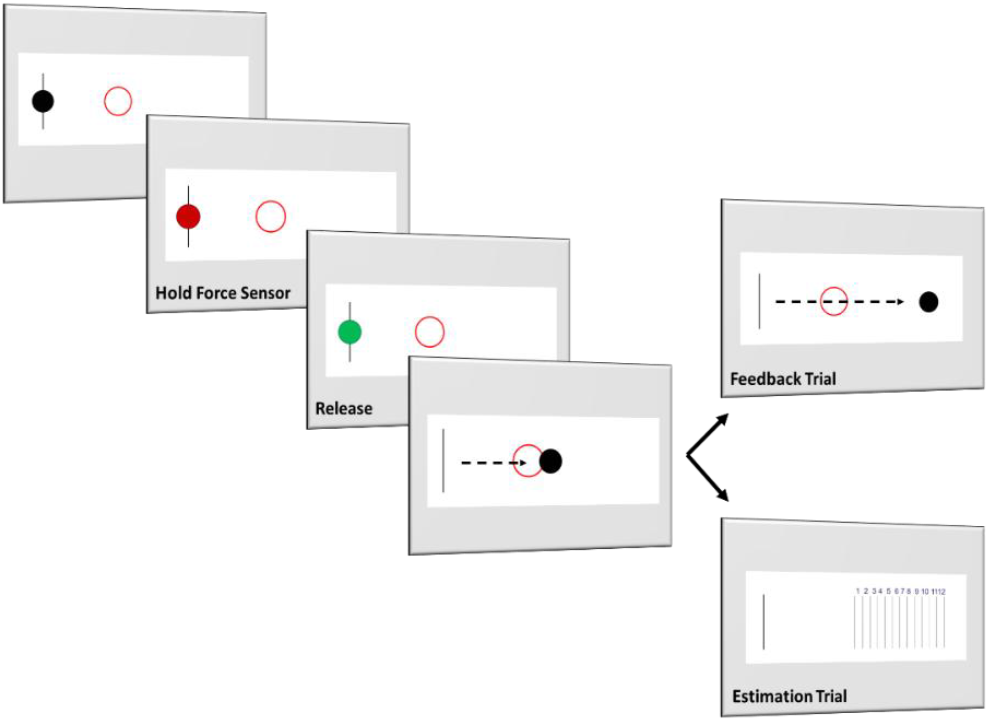
Pipeline of the ‘Goal Prior Assay’ task originated by Hezemans et al (2020) as a behavioural paradigm of apathy under a Bayesian conceptual framework. When the ball appears, participants are asked to press a force sensor, at which point the ball will turn red. After 3 seconds the ball turns green, and participants are instructed to release the force sensor. At this point the virtual ball travels across the screen, with the distance travelled determined by the force applied to the sensor within the initial 3-second window. In non-catch trials, the ball is seen to stop. In pseudo-random catch trials, the ball disappears early in its trajectory and participants are asked to estimate the final position of the ball using an estimation scale that appears on screen.

During the experiment, saccades were recorded using a long-range mounted EyeLink 1000 (SR User Manual, v1.5.2). Standard 9-point calibration and validation was used to calibrate the eye tracker at the beginning of the task.

In the second testing session, participants completed 7T MRI on a Siemens MAGNETOM Terra scanner to acquire structural data. Following the scan, participants completed a neuropsychological battery including the Revised Addenbrookes Cognitive Examination (ACER)^63^, Frontal Assessment Battery (FAB)^64^, INECO Frontal Screening (IFS)^65^ and Hayling sentence completion task^66^.

### 4.4 Imaging Data Collection

MEG data was acquired at a frequency of 1kHz in a magnetically shielded room using the 306-channel MEGIN TRIUX neuro system which includes 204 planar gradiometers and 102 magnetometers. Eye movements were measured using pairs of vertical and horizontal EOG electrodes, and heart rate was similarly measured with a pair of ECG electrodes placed on the wrists. Prior to data acquisition, a 3D digitizer (Fastrak Polyhemis Inc.) was used to record the following: 1) three fiducial points, one on the nasion and one each on the left/ right pre-auricular points, 2) the location of 5 HPI coils, and 3) ∼300 ‘head points’ across the scalp.

All MEG preprocessing was conducted in MATLAB v2018a using the Statistical Parametric Mapping (SPM12) toolbox based on a pipeline established by Vaghari et al (2022)^67^. Only non-catch trials were used in MEG analysis. The raw MEG data was first preprocessed using Maxfilter software (version 2.2.14, Neuro-TRIUX), including standard signal-space separation, movement compensation and translation to default space. Epochs of 2s [-1000 1000] were extracted from the data, time-locked around one of three task events: force sensor press, force sensor release, and ball stop. Data was filtered (0.5-150Hz bandpass filter followed by 48-52Hz notch filter) and ocular and cardiographic artefacts were flagged using independent components analysis^68^. Further artefact rejection was conducted using software from the OHBA Software Library. Finally, the cleaned data was manually coregistered using the fiducials digitised at MEG acquisition and the denoised 7T MRI scan for each participant. A 7T template^69^ was used in cases where a scan was unavailable (N=2).

MR data was acquired using a MAGNETOM Terra scanner (Siemens) with 32-channel head coil at the Wolfson Brain Imaging Centre. For the purposes of MEG coregistration, we acquired a high resolution isotropic whole brain Magnetization Prepared 2 Rapid Acquisition Gradient Echoes (MP2RAGE) sequence (repetition time=4300 ms, echo time=1.99 ms, resolution=99 ms, bandwidth=250 Hz/px, voxel size=0.75 mm3, field of view= 240 × 240 × 157mm, acceleration factor (A ≫ P) =3, flip-angle=5/6° and inversion times=840/2370 ms).

## 5. Data Analysis

For both behavioural analyses and network modelling we primarily used Bayesian statistics, which were corroborated with secondary frequentist statistics where appropriate. The Bayes Factor (B) was interpreted in keeping with established conventions ^70,71^. Standard evidentiary thresholds were used for moderate (>3), strong (>10) or very strong (>100)^72^ evidence in favour of the alternate hypothesis, or 1/3, 1/10 and 1/100 for corresponding strength of evidence for the null. Code for this data analysis pipeline are available at https://github.com/BeccaSue99/Pbl-MEG.

### 5.1 Regression Analyses

Behavioural analyses were conducted with custom scripts in RStudio v2023, using R v4.2.2.

#### Hypothesis 1

There is a negative correlation between trait apathy and prior precision.

Hypothesis 1 is a replication of previous work by Hezemans et al. (2020) and is assessed using a Bayesian correlation. Prior precision was calculated as the regression slope of estimation error on performance error for each participant. We expect participants with high levels of prior precision to be systematically and more strongly biased in the direction of the target. This systematic bias results in a tendency to overestimate the final position of the virtual ball when it undershoots the target, and underestimate when the ball overshoots the target. For this reason, the gradient of the regression slope can be used as a proxy of prior precision.

Trait apathy was based on the Apathy Motivation Index score. Correlations were calculated between prior precision and other self-report measures to explore potential links with impulsivity and anhedonia.

### 5.2 Bayesian Hierarchical Modelling

In order to confirm the target as the location of the prior we implemented Bayesian Hierarchical Modelling (BHM) as detailed in Hezemans et al (2020).

For any given catch trial we consider the sensory evidence to be centred on the true location of the ball, and the posterior to be centred on the participant’s estimation, multiplied by the pixel width of each estimation bin in order to have both measures in a comparable range. We tested three potential models of the prior each of which had unknown variance: (i) the prior is centred on the target, (ii) the prior is centred on the target with a set perceptual shift, and (iii) the prior is centred on the true performance of the participant. Models used Markov chain Monte Carlo sampling to approximate the posterior distributions of each parameter. For each model, we used eight independent chains with 2,000 samples, discarding the first 1,000 samples as the “burn-in” period. We compared the models using the Widely Applicable Information Criterion (WAIC).

#### Hypothesis 2

There is a difference in prior precision **(A)** between effort blocks, but **(B)** not between rewarded and unrewarded blocks.

We used BHM to investigate hypothesis 2 which sought to replicate additional findings from Hezemans et al. (2020).

### 5.3 Eye Tracking

Eye tracking was analysed using custom scripts generated in MATLAB v2018a and RStudio v2023, using R v4.2.2.

#### Hypothesis 3

Saccadic eye movements establish an implicit measure of prior precision in the ‘Goal Priors Assay’ task.

The focus of the eye tracking analysis was the sample immediately preceding the appearance of the estimation grid onscreen. Samples in which gaze along the y position was more than two standard deviations from the mean were removed from further analysis due to concerns that the participant may not have been focussed on the stimulus screen.

The quality of the eye tracking data was checked by first correlating gaze position along the x axis with the final location of the ball in non-catch trials. Following this we replicated the regression analysis and BHM detailed above, replacing explicit estimations with gaze x position at the time sample immediately preceding the appearance of the estimation grid.

### 5.4 Source Reconstruction

Neurophysiological analyses were conducted with custom scripts in MATLAB v2018a using the SPM12 toolbox.

Following coregistration, we implemented a beamformer using gradiometers only to localise sources as we intended to use this localization to inform selection of a network for later modelling using gradiometers only. 250ms thresholded contrasts (p[uncorrected] < 0.001, k=100) were generated from the beamformed data using the 1 second epoch around which the ball stops. For each image the baseline was selected as the 250ms time window immediately before, such that significant activity reflected *changes* in neural activity across the epoch.

Our preregistration specified a potential network for this task with sources in the right inferior frontal gyrus, left premotor and left motor cortices, using coordinates from Hughes et al. (2018). However, we anticipated that the exact coordinates of these nodes may change depending on results from source reconstruction.

As seen in *figure 7*, there was a bilateral increase in power in the motor and premotor cortex in the 500ms across ball stop. There was also a significant change in power in the right inferior frontal gyrus. There was a bilateral decrease in power across the frontopolar cortex and the anterior inferior frontal gyrus, between 250ms to 500ms after ball stop.

**Figure 7.**
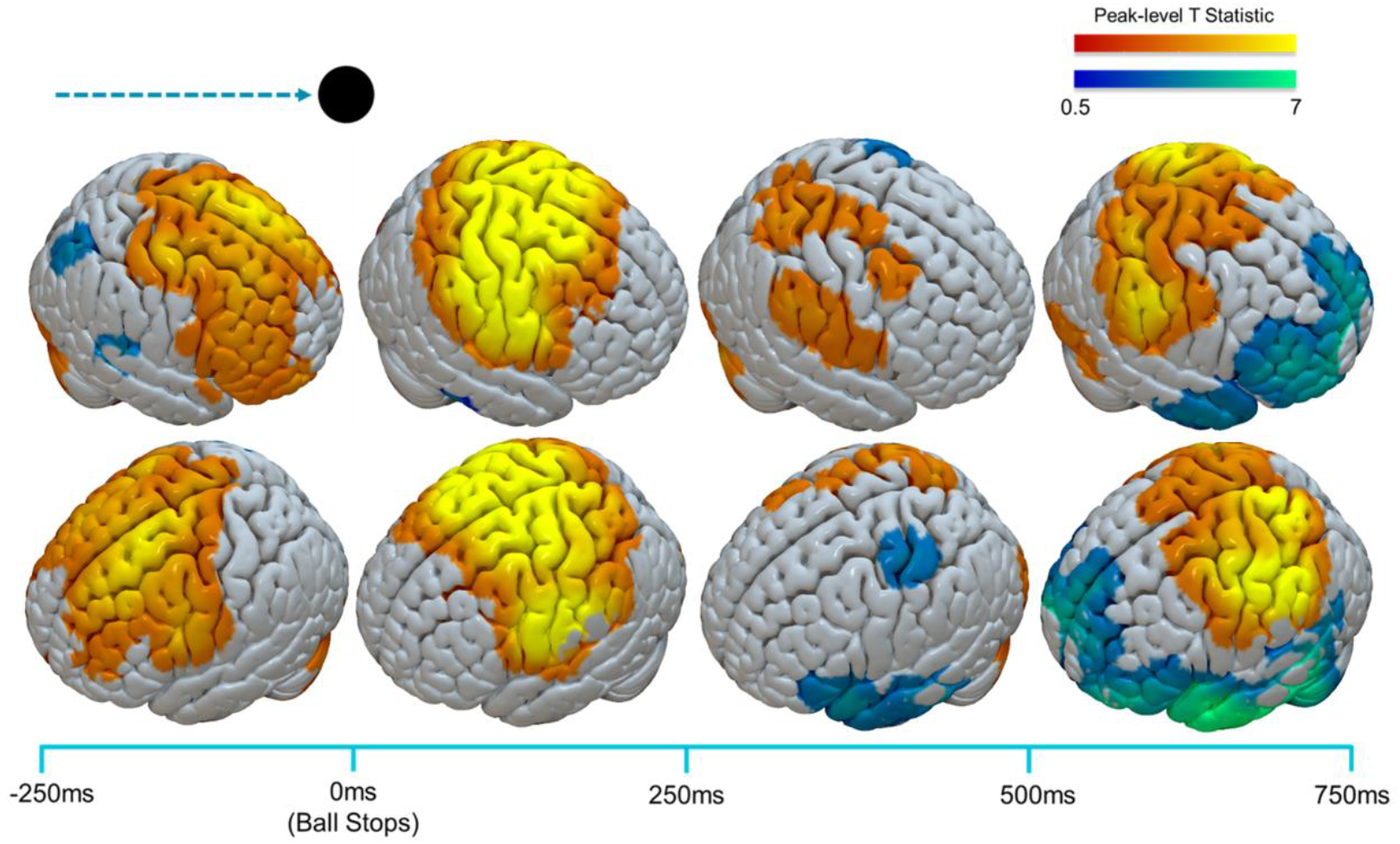
Reconstructed sources using LCMV beamformer with gradiometers only in the one second around ball stop in non-catch trials. Clusters thresholded at p[uncorrected] < 0.001, k=100.

Given significant activity was evident in all three preregistered sources, we selected our coordinates for future analysis as the nearest local peak in activity to the preregistered coordinates: right inferior frontal gyrus (pars orbitalis) [56; 26; -12], left premotor cortex [-14; 16; 58], and left motor cortex [-37; -25; 64]. In the case of the motor coordinate, due to widespread activity, we used the original coordinate from Hughes et al (2018).

### 5.5 Spectral Analysis

Following the identification of time windows, frequency ranges and coordinates using beamforming, we extracted local field potentials from gradiometers only at each of the selected coordinates.

#### Hypothesis 4

Participants with low prior precision have a reduction in beta band oscillations in the prefrontal cortex.

In order to test hypothesis 4, that there is a link between beta activity and apathy scores, we calculated the relative power spectral density (PSD) of LFP’s at each coordinate, but focussing particularly on the prefrontal cortex as previously specified. Correlations between relative beta power and apathy were calculated using custom scripts in RStudio v2023, with beta power calculated as mean power across each time window in the 13-30Hz range, as there was not a clear beta peak in the spectral data (see *figure 8*). We also ran analyses with the beta range split into low (13-22Hz) and high (22-30Hz) beta ranges.

**Figure 8.**
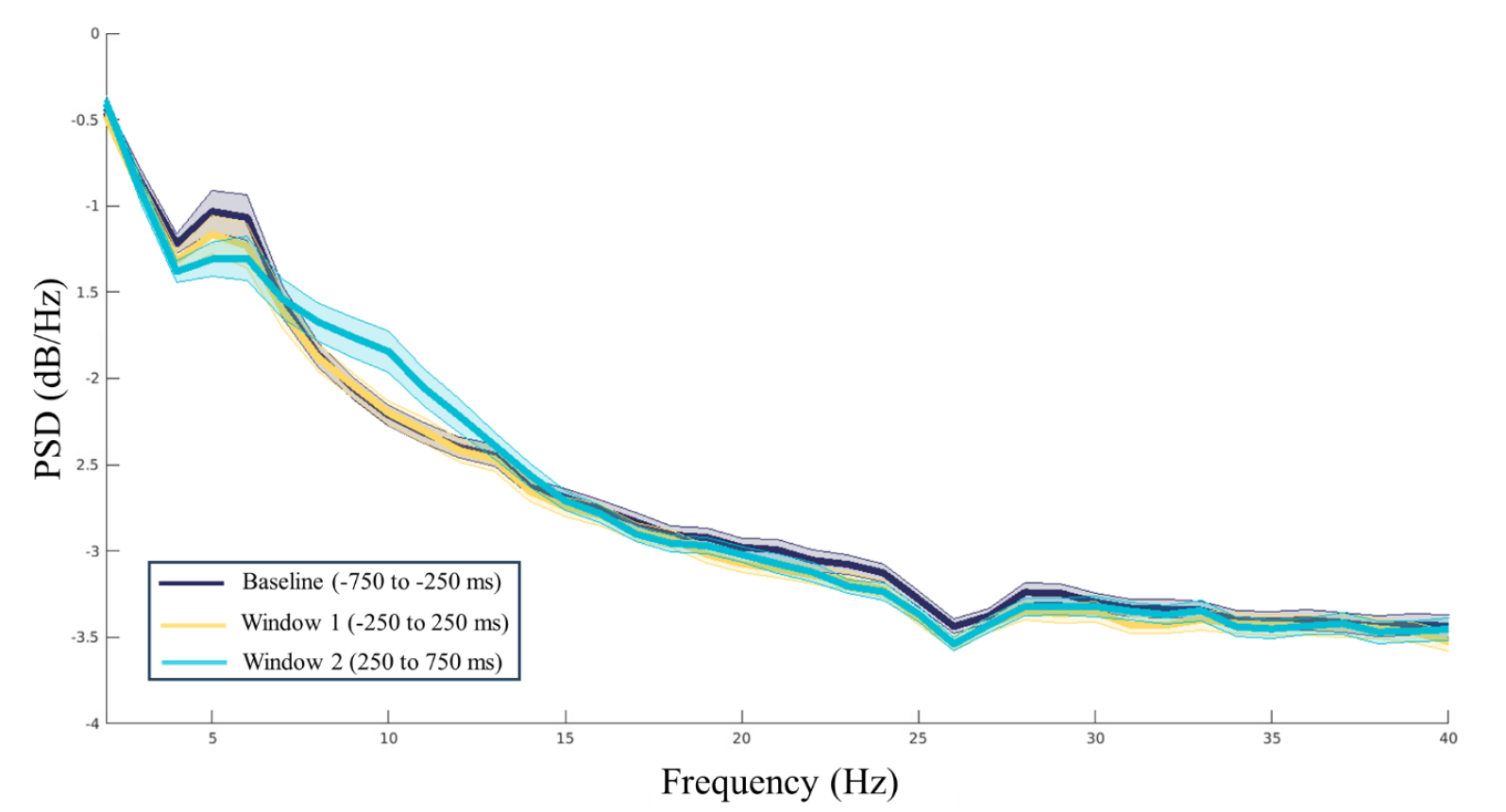
Relative power spectral density curve calculated from the gradiometer-only local field potential extracted from the right inferior frontal gyrus at baseline [-750ms -250ms], time window 1 [-250ms 250ms] and time window 2 [250ms 750ms]. 0ms represents the time at which the ball stops.

### 5.6 Dynamic Causal Modelling

#### Hypothesis 5

The inclusion of prior precision as an empirical prior of the generators for physiological responses can improve model fidelity in the prefrontal and motor cortices.

Our first-level analysis used a conductance-based dynamic causal model (DCM; Variant CMM-NMDA) of cross spectral densities to model the extracted local field potentials of the 500ms around ball stop. We checked the quality of model fit by calculating both variances explained and the correlation between observed and predicted response.

We then compared evidence associated with two models within this framework at a group level using Parametric Empirical Bayes (PEB)^73^. The first PEB model included prior precision as an empirical prior of synaptic physiology, whereas the second featured no empirical priors related to task performance. Hypothesis 5 was assessed by comparing respective model evidence using Bayesian evidentiary thresholds as previously specified. Given our hypotheses concerning GABAergic contributions to top-down communication, we particularly focussed our interpretation of parameters on the synaptic gain of the superficial pyramidal neurons and time constants of GABA. Effects were considered significant if the posterior probability exceeded 95%.

## 6. Acknowledgements

This work has been funded by the Wellcome Trust (220258), the Cambridge Trust, the Medical Research Council (MC_UU_00030/14; MR/T033371/1), the NIHR Cambridge Biomedical Research Centre (NIHR203312), the Cambridge Centre for Parkinson-plus and the Holt fellowship Cambridge Home and EU Scholarship Scheme, James F. McDonnell Foundation, and Evelyn Trust. The views expressed are those of the authors and not necessarily those of the NIHR or the Department of Health and Social Care. For the purpose of open access, the authors have applied a CC BY public copyright licence to any Author Accepted Manuscript version arising from this submission.

